# *mtor* Haploinsufficiency Ameliorates Renal Cyst Formation in Adult Zebrafish *tmem67* Mutants

**DOI:** 10.1101/2019.12.20.883710

**Authors:** Ping Zhu, Qi Qiu, Peter C. Harris, Xiaolei Xu, Xueying Lin

## Abstract

Although zebrafish embryos have been utilized to study ciliogenesis and to model polycystic kidney disease (PKD), adult zebrafish remain unexplored. Here, we report the generation and characterization of a zebrafish mutant of *tmem67*, a homologue of the mammalian causative gene for Meckel syndrome type 3 (MKS3). Although a small population of homozygous embryos exhibited pronephric cysts, all mutants were able to survive to adulthood and developed progressive mesonephric cysts with full penetrance. The cysts in the adult zebrafish kidneys manifested features of mammalian PKD, including switching of cyst origin from the proximal tubules to the collecting ducts, increased proliferation of cyst-lining epithelial cells, and hyperactive mTOR signaling. Consistent ciliary abnormalities were observed in both the embryonic and adult zebrafish mutants compared with the wild-type fish, including shorter and fewer single cilia in the distal pronephros and all segments of the mesonephros and greater numbers of multiciliated cells (MCCs). Lack of single cilium preceded cystogenesis, suggestive of a primary defect. Expansion of MCCs occurred after pronephric cyst formation and was inversely correlated with the severity of cystogenesis in young adult zebrafish, suggesting an adaptive action. Interestingly, mTOR inhibition ameliorated renal cysts in both the embryonic and adult zebrafish models; however, it only rescued ciliary abnormalities in the adult mutants. In summary, we have established a *tmem67* mutant as the first adult zebrafish PKD model, revealed a novel aspect of cilium regulation, and identified sustained mTOR inhibition as a candidate therapeutic strategy for *tmem67*-based PKD.

**Significance Statement:** While zebrafish embryos are well recognized for their value in studying ciliogenesis and polycystic kidney disease (PKD), adult zebrafish have not commonly been used. Here, we report the establishment of the first adult zebrafish model for PKD, which exhibits characteristics of mammalian PKD and shows kidney ciliary abnormalities consistent with those observed in an embryonic model. We also provide evidence for mTOR inhibition as a therapeutic strategy for this particular type of cystogenesis. Compared to the embryonic model, the adult fish model exhibits a spectrum of progressive pathogeneses and enables ciliary abnormalities to be discerned as either primary or secondary to cystogenesis. We believe that this novel adult fish model will facilitate mechanistic studies and therapeutic development for PKD.

## Introduction

*TMEM67* is a major causative gene for Meckel syndrome (MKS; the gene was formerly named *MKS3*), accounting for approximately 15% of MKS cases; it is also a causative gene for Joubert syndrome (JBS).^1–4^ MKS is an autosomal recessive and perinatal lethal disorder that is characterized by a spectrum of abnormalities, including renal cysts in more than 95% of patients, central nervous system defects such as encephalocele, liver fibrosis, and sometimes polydactyly.^5–7^ Spontaneous animal models have been used to study MKS3, including the *wpk* rat carrying a naturally occurring single point mutation in the *Tmem67* gene, the *bpck* mouse with the causative *Tmem67* gene encompassed in a large 245-kb deletion, and even a sheep model.^1, 8–10^ The murine models capture some characteristics of MKS3 patients, such as polycystic kidneys and hydrocephalus, but not encephalocele, biliary abnormalities, polydactyly, while the sheep has the characteristic hepatorenal disease.^8–12^ A targeted knockout mouse (*Tmem67*^*tm1*(*Dgen*/*H*)^) has also been generated that exhibits pulmonary hypoplasia, cardiac malformation, kidney cysts, and some encephalocele and dies by postnatal day 1 (P1) due to pulmonary and/or cardiac defects.^13–15^ Of note, an effective therapy for MKS has not yet been reported.

*TMEM67* encodes a transmembrane protein (meckelin) that is localized in the ciliary transition zone of the renal epithelium;^1, 13, 16^ thus, *TMEM67*-based polycystic kidney disease (PKD) is considered a ciliopathy. However, a range of ciliary phenotypes has been noted among different MKS3 models. For instance, *bpck* mice, *wpk* rats, and human MKS3 fetuses have elongated renal cilia; murine *Tmem67* mutants exhibit shorter and fewer cilia than wild-type mice; and ovine *TMEM67* mutants display both very long and very short cilia in their cystic kidneys.^9–11, 13^ Moreover, compared with those from wild-type mice, MEFs derived from *Tmem67* knockout mice present longer, normal, or no cilia; similar findings have been obtained with *Tmem67* shRNA-expressing cells.^11, 13, 14, 17^ Therefore, additional studies are needed to define ciliary defects in MKS3 models and to clarify the relationship between ciliary defects and cyst development.

Embryonic zebrafish have long been used to study PKD owing to their transparency and the efficiency of genetic manipulation.^18–21^ Embryonic zebrafish have also proven to be excellent models for *in vivo* analyses of ciliogenesis and cilium maintenance.^22^ The cilia in the zebrafish pronephric kidney include single motile cilia arising from the majority of epithelial cells and cilia arranged in multicilia bundles existing in the proximal straight tubule (PST) and distal early (DE) segment.^23–26^ Single-ciliated cells (SCCs) and multiciliated cells (MCCs) form an intercalated ‘salt and pepper” pattern in the PST-DE region that is controlled by Notch signaling.^25, 26^

Because the pathogenesis of PKD cannot be fully recapitulated during 8 days of embryogenesis, we turned our attention to adult zebrafish. The adult zebrafish mesonephric kidney undergoes similar branching morphogenesis and segment organization (proximal tubule (PT), distal tubule (DT), and collecting duct (CD)) as the mammalian kidney, although it contains much fewer nephrons (~200 vs 1 million in humans) and lacks the loop of Henle. Because the loop of Henle functions to preserve water in mammals, it is unnecessary for freshwater zebrafish to have this segment.^27–30^ Whether adult zebrafish can be used to model PKD and to develop therapies has not been explored.

Here, we report the generation of a zebrafish *tmem67* mutant and the characterization of its renal and ciliary phenotypes. We noted pronephric cysts in the embryos and mesonephric cysts in the adult fish; thus, we have established the first adult zebrafish model of PKD. We defined ciliary abnormalities during embryogenesis and adulthood and correlated these defects with cyst development. Finally, we revealed that mTOR inhibition is a candidate therapeutic strategy for Tmem67-associated renal cyst formation.

## Methods

### Zebrafish strains

Zebrafish (WIK) were maintained under standard laboratory conditions, and experiments were carried out in accordance with the policies of the Mayo Clinic Institutional Animal Care and Use Committee. A hypomorphic *mtor* strain was identified from our insertional mutagenesis screen.^31^

Zebrafish *tmem67* mutants were generated using the Golden Gate TALEN assembly protocol and library.^21, 32–34^ TALEN mRNAs were injected into embryos at the one-cell stage. F1 adults carrying germline mutations were identified by PCR (forward primer: 5′-TGTATAGGACTGGCATGTGAG-3′; reverse primer: 5′-AGGGATTGCCATTCCCATC-3′) and XmnI restriction enzyme digestion. Frameshift mutations were identified by sequencing, and the zebrafish were further outcrossed to reduce potential off-target effects. All experimental fish used in this study were F3 or F4 animals.

### In situ hybridization

Whole-mount *in situ* hybridization was performed as previously described using riboprobes that were generated from T7 promoter sequence-tagged PCR products.^35^

### Histological analysis

Zebrafish embryos were analyzed by hematoxylin and eosin (HE) staining as we have described previously.^21^ Adult zebrafish kidneys were also collected as described in another study.^36^ Briefly, each fish was euthanized with 0.2% Tricaine. All of its internal organs were removed except the kidney, which was attached to the dorsal wall of the abdominal cavity. The fish body with the kidney was then fixed in 4% paraformaldehyde (PFA) overnight at 4°C. On the next day, the kidney was carefully detached from the abdominal wall and subjected to paraffin embedding and HE staining. If a tubule was dilated to an area exceeding 0.03% of the total kidney area, it was considered a cyst. The areas of renal cysts as percentages of the total tissue area were calculated using ImageJ software.

### Immunofluorescence labeling

Cilia in zebrafish embryos were visualized by whole-mount immunofluorescence staining using antibodies against acetylated α-tubulin (Sigma-Aldrich), as previously described.^37, 38^ Antibodies against α6F (Developmental Studies Hybridoma Bank) were included for staining of the pronephros.

Immunofluorescence analysis of adult zebrafish kidneys was performed on cryo-sections as described previously.^21, 28, 29^ The kidneys were dissected, fixed in 4% PFA/0.1% DMSO overnight at 4°C and then permeabilized with 5% sucrose for 30 minute followed by 30% sucrose overnight. The following day, the kidneys were embedded in tissue freezing medium (Electron Microscopy Sciences) and cryo-cut at 10 μm (Leica CM3050S). Immunofluorescence analysis of adult zebrafish kidneys was also conducted on paraffin sections as described previously.^39^ Renal tubular segments were labeled with alkaline phosphatase (AP, Invitrogen), rhodamine *Dolichos biflorus* agglutinin (DBA), and *Lotus tetragonolobus* lectin (LTL) (Vector Laboratories). Nuclei were labeled with SYTOX, DAPI (Vector Laboratories), or propidium iodide (PI) (Sigma-Aldrich). Anti-PCNA and anti-acetylated α-tubulin antibodies (Sigma-Aldrich) were used for immunostaining. Images were acquired using a Zeiss Axioplan II microscope equipped with ApoTome and AxioVision software (Carl Zeiss Microscopy).

### Ultrastructural analysis

Adult zebrafish kidneys were fixed in Trump fixative (4% PFA, 1% glutaraldehyde). The remaining procedures for transmission electron microscopy (TEM) and scanning electron microscopy (SEM) were performed according to standard methods at the Electron Microscopy Core Facility of the Mayo Clinic in Rochester, Minnesota.

### Western blotting

Adult zebrafish kidneys were homogenized in RIPA lysis buffer (Sigma-Aldrich) as described previously.^21^ The following antibodies were used for western blotting: anti-S6 ribosomal protein (Cell Signaling Technology), anti-phospho-S6 ribosomal protein (Ser240/244) (Cell Signaling Technology), and anti-actin (Sigma-Aldrich).

### Rapamycin treatment of adult zebrafish

Rapamycin (LC Laboratories) was administered at the specified dose to adult zebrafish via oral gavage as described previously.^40^ Treatment was conducted daily and lasted for one month.

### Statistical analysis

The data are presented as the mean ± s.d. Comparisons between two groups were performed with two-tailed Student’s *t* tests, and a *P* value <0.05 was considered to indicate significance.

## Results

### *tmem67*^*e3/e3*^ embryos exhibit pronephric cysts with partial penetrance

*tmem67*, the only zebrafish homologue of mammalian *TMEM67*, is located on chromosome 16 (Figure 1A).^12^ Its transcript was detected in the kidney, neural tube, otic vesicle, brain, and retina during embryogenesis (Figure 1B) and in all major organs of adult fish (Supplemental Figure 1). To investigate the role of TMEM67 in renal cyst formation and ciliogenesis, we designed TALEN pairs targeting exon 3 of *tmem67*, because pathogenic mutations are mostly found in the exons encoding the N-terminal domains of meckelin.^3, 41, 42^ We analyzed two *tmem67* mutant alleles that resulted in a premature stop codon, including the M1 allele containing a 1-bp insertion and the M2 allele containing a 5-bp deletion (Figure 1A,C). Because both alleles exhibited the same phenotypes, we have presented data only from the M1 allele and have renamed this allele *tmem67^e3^*.

**Figure 1.**
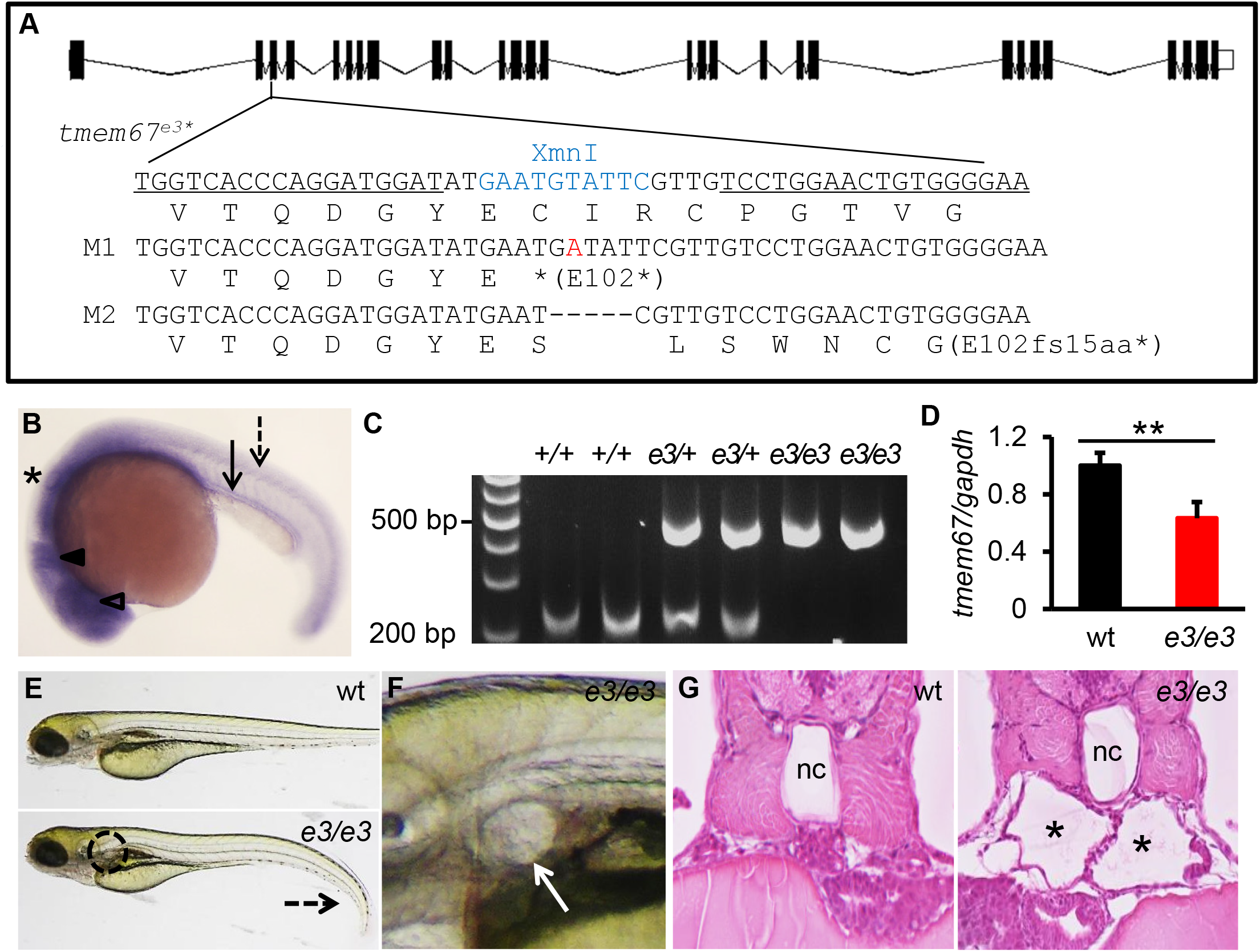
Zebrafish *tmem67*^*e3/e3*^ embryos develop partially penetrant pronephric cysts. (**A**) Schematic diagram of the exon-intron structure and two mutant alleles. The TALEN recognition sequences are underlined. Mutations in both the M1 and M2 alleles cause a coding frameshift and premature stop codon (*). Additions are indicated in red, and deletions are indicated with dashes. The restriction enzyme recognition site is highlighted in blue. The corresponding amino acid sequence is shown below the DNA sequence. (**B**) *In situ* hybridization showing *tmem67* expression in an 18-somite embryo. The pronephric kidney is indicated with an arrow, the neural tube with a dashed arrow, the otic vesicle with asterisks, the brain with a closed arrowhead, and the lens with an open arrowhead. (**C**) Genotyping of *tmem67*^*e3/e3*^ embryos. PCR products amplified from wild-type alleles could be digested by XmnI to produce 257-bp and 237-bp fragments, while PCR products amplified from mutated alleles could not be cleaved. (**D**) Quantitative PCR analysis of *tmem67* transcripts. Embryos at 5 dpf were subjected to RNA extraction, cDNA synthesis, and quantitative PCR analysis. The results were normalized to those for *gapdh*. The data are presented as the mean ± s.d. from three independent experiments. Eight to ten embryos per genotype were examined. **: *P*<0.01. (**E**) Gross morphology of *tmem67*^*e3/e3*^ embryos. Mutants developed ventral curvature of the body (dashed arrow) and pronephric cysts (dashed circle). Shown are embryos at 4 dpf. (**F**) Enlarged view of the pronephric cyst in E. (**G**) The glomerular neck region of the pronephros was dilated in some *tmem67*^*e/e3*^ embryos (asterisks). HE staining of JB-4 sections in day 3 embryos is shown. nc: notochord.

In *tmem67*^*e3/e3*^ embryos, *tmem67* transcript levels were reduced by nearly 40%, likely due to nonsense-mediated mRNA decay (Figure 1D). We could not quantify meckelin levels due to the lack of antibodies recognizing the zebrafish protein. Approximately 40% of the *tmem67*^*e3/e3*^ embryos exhibited ventral body curvature, and 20% developed pronephric cysts; furthermore, cystic embryos were more likely to be curved than non-cystic embryos (Figure 1E-G). Unlike rodent knockout animals and zebrafish morphants, the *tmem67*^*e3/e3*^ mutants did not exhibit apparent defects in convergence extension or neural tube development and rarely developed hydrocephalus (Supplemental Figure 2 and data not shown).^8, 9, 12, 14^ We assessed the possibility that the mutations were hypomorphic but we did not note any skipping of the targeted exon via alternative splicing events (Supplemental Figure 3).

### *tmem67*^*e3/e3*^ embryos progressively lose single cilia in the pronephros

Next, we assessed renal cilia. We noted significantly shorter cilia in both the proximal and distal pronephron segments in *tmem67*^*e3/e3*^ embryos compared to wild-type at 26 hours post fertilization (hpf) (Figure 2A,F), when developing pronephros do not yet have filtration functions and when multicilia clusters are not yet present.^18, 25, 26^ Because pronephric cysts are visually detectable at 2 days post fertilization (dpf),^18^ manifesting as dilations in the glomerulus-neck region, we grouped the mutants into those with cysts (designated as +EC) and without cysts (designated as −EC). Compared with wild-type organisms, both +EC and −EC mutants had significantly shorter and fewer single cilia at the segments caudal to the multiciliated region (Figure 2B,E,G,H) and had less but only marginally shorter single cilia at the segments rostral to the multiciliated region (Figure 2B,C,G). The overall reductions in both cilium length and number were comparable between +EC and −EC embryos (Figure 2G-H). In contrast, multicilia bundles and MCCs, as indicated by *odf3b* expression, remained normal at this stage (Figure 2B,D,I-K). At 4 dpf, distal single cilia were absent from all mutants, but the number of multicilia bundles continued to be normal; however, the numbers of cells with MCC fates were increased in the +EC embryos, suggesting that the number of cilia bundles may increase at a later time (Supplemental Figure 4). In contrast to the pronephros, other tissues, such as Kupffer’s vesicle and the neural tube, had unaffected cilia (Supplemental Figure 5).

**Figure 2.**
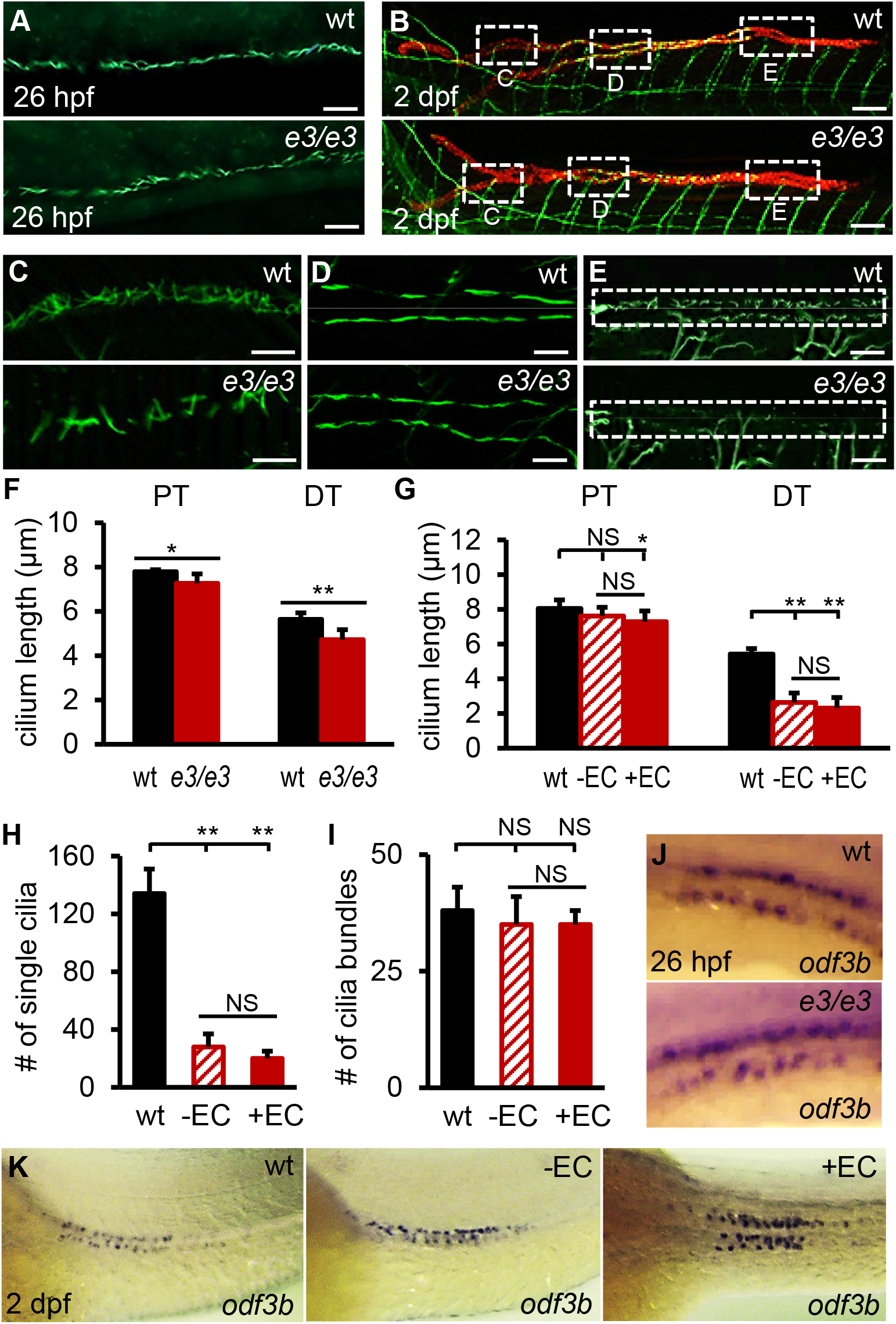
*tmem67*^*e3/e3*^ embryos have shorter and fewer single cilia in the distal pronephros than wild-type embryos. (**A**) Pronephric cilia in 26 hpf embryos were examined by whole-mount immunostaining using an α-acetylated tubulin antibody. (**B**) Pronephric cilia from a wild-type embryo and a mutant without visually detectable pronephric cysts at 2 dpf were illustrated by coimmunostaining using antibodies against α-acetylated tubulin (green) and Na^+^/K^+^ ATPase (α6F, red; labels the pronephros). The boxed areas are enlarged in C-E. (**C-E**) Enlarged representative images of α-acetylated tubulin staining. Proximal single cilia are shown in C, multicilia bundles in D, and distal single cilia in E. (**F**) Quantification of renal cilium length in 26 hpf embryos. The cilia at the proximal end (PT) and distal end (DT) of the pronephros were measured. (**G**) Quantification of renal single-cilium length in 2 dpf embryos. Single cilia in the tubular segment immediately rostral to the multiciliated region (PT) or caudal to the multiciliated region (DT) were measured. (**H**) The total numbers of single cilia in the distal tubules of both pronephros were counted at 2 dpf. (**I**) Quantification of multicilia bundles at 2 dpf. (**J**,**K**) MCCs were examined by *in situ* hybridization using an *odf3b* riboprobe. Shown are representative embryos at 26 hpf (J) and 2 dpf (K). The data are presented as the mean ± s.d. from two (F) or four (G-I) independent experiments. A total of 16-22 embryos per group per age were examined. *: *P*<0.05; **: *P*<0.01; NS: *P*>0.05, not statistically significant. Scale bars: 20 μm (A,C-E) and 50 μm (B).

The +EC embryos showed failure of anterior migration and convolution of tubular epithelial cells (Supplemental Figure 6A). Consistently, these embryos had a condensed MCC domain and shortened PST and DE segments (Figure 2K, Supplemental Figures 4D and 6B,C). Because anterior migration and convolution require fluid flow in the tubule,^24, 43–45^ these data suggest impaired renal fluid movement in the +EC embryos, which could contribute to pronephric cyst formation.

### Adult *tmem67*^*e3/e3*^ fish develop progressive mesonephric cysts

Most *tmem67*^*e3/e3*^ embryos, including cystic ones, survived for at least 15 months, with approximately 40% of fish exhibiting wavy bodies (Figure 3A). Independent of body shape, the mutant kidneys were significantly enlarged (Figure 3B,C), and the renal tubules were progressively dilated in male fish and, to a much less extent, female fish (Figure 3D,E and data not shown). Because the renal phenotype was milder than that in rodent models, we examined the expression of other MKS genes. While *tmem67* transcript levels were reduced by approximately 50%, *mks1*, *cep290*, and *cc2d2a* were activated in the mutant kidney, suggesting genetic compensation that has been associated with TALEN-mediated mutagenesis (Supplemental Figure 7).^46–48^ Biliary abnormalities were not observed in the *tmem67* fish up to 12 months.

**Figure 3.**
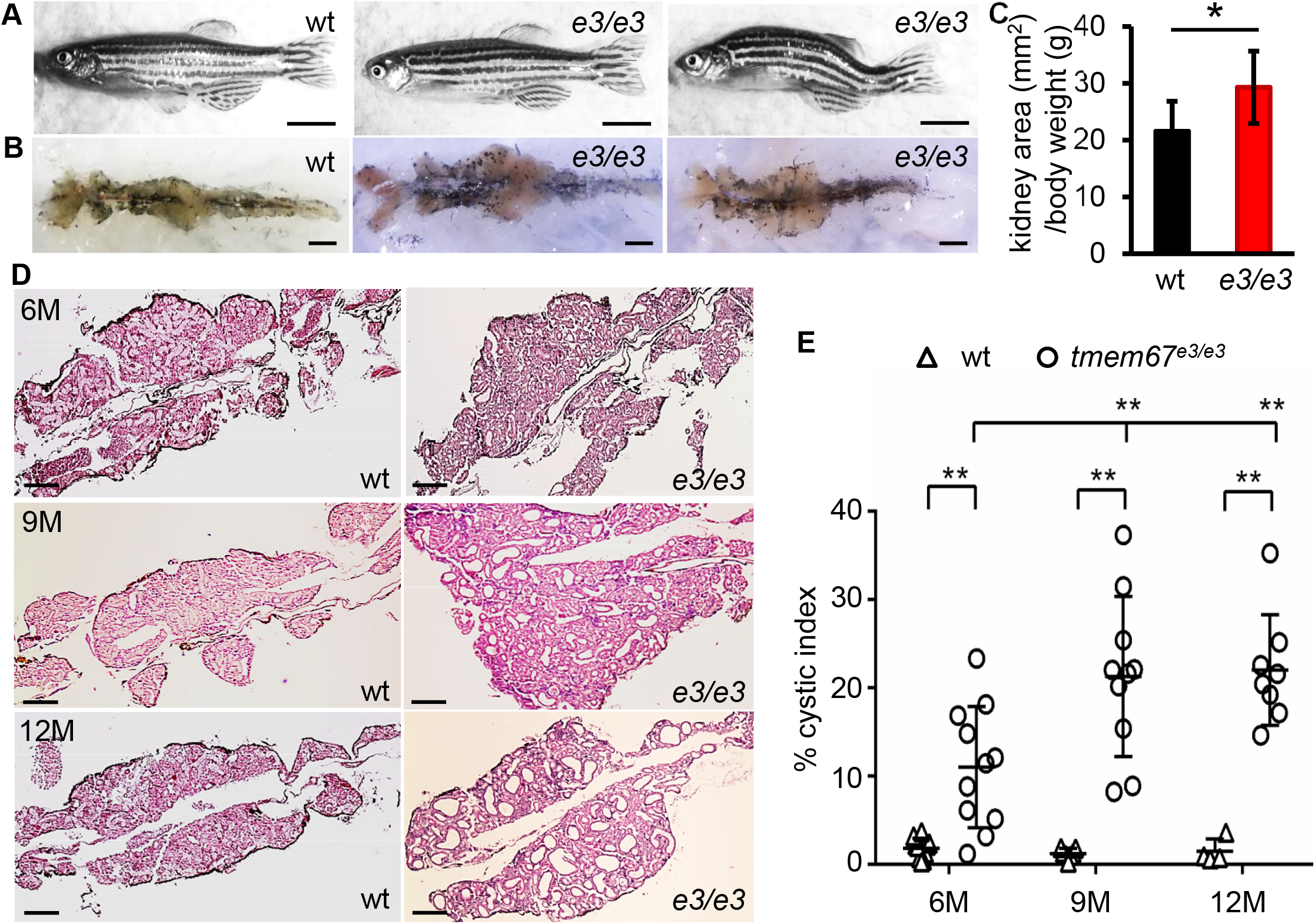
Adult *tmem67*^*e3/e3*^ fish develop progressive renal cysts. (**A**) *tmem67*^*e3/e3*^ fish were either morphologically indistinguishable from their wild-type siblings or curved in shape. Shown are 12-month-old fish. (**B,C**) *tmem67*^*e3/e3*^ fish had larger kidneys than wild-type fish. Twelve-month-old fish were fixed in 4% PFA, and then the kidneys were dissected for size measurement (B). The kidney area was normalized by the body weight (C). Eight male fish from each group were analyzed. (**D,E**) *tmem67*^*e3/e3*^ fish exhibited renal cysts. Kidneys were collected at the indicated ages, and HE staining was performed on paraffin sections. Representative images (D) are shown. The cyst area/total kidney area (cystic index) was calculated (E). Four to 11 male fish from each group at each time point and 3 sections per kidney were analyzed. The data are presented as the mean ± s.d. *: *P*<0.05; **: *P*<0.01 (C,E). Scale bars: 7 mm (A), 1 mm (B), and 100 μm (D).

In rodent *Tmem67* models, renal cysts present as a mixture of PT dilation and CD enlargement in newborns and are mainly restricted to the CD at the later stage.^8, 9^ A similar progression was noted in the *tmem67*^*e3/e3*^ zebrafish. The proportion of PT cysts, marked by AP, decreased from 85% of all cysts to 33% from 6 to 12 months; DT cysts, labeled by DBA, slightly increased; and AP/DBA(−) cysts, presumptively a mixture of CDs and tubules with dedifferentiated epithelial cells, significantly increased (Figure 4A,B). We further costained samples with DBA and LTL, which labels PTs as well as the major CDs in the zebrafish kidney,^28, 29^ and found that the numbers of LTL cysts only moderately reduced and the numbers of unstained cysts moderately increased with time, indicating more CD cysts at a later stage (Figure 4C,D).

**Figure 4.**
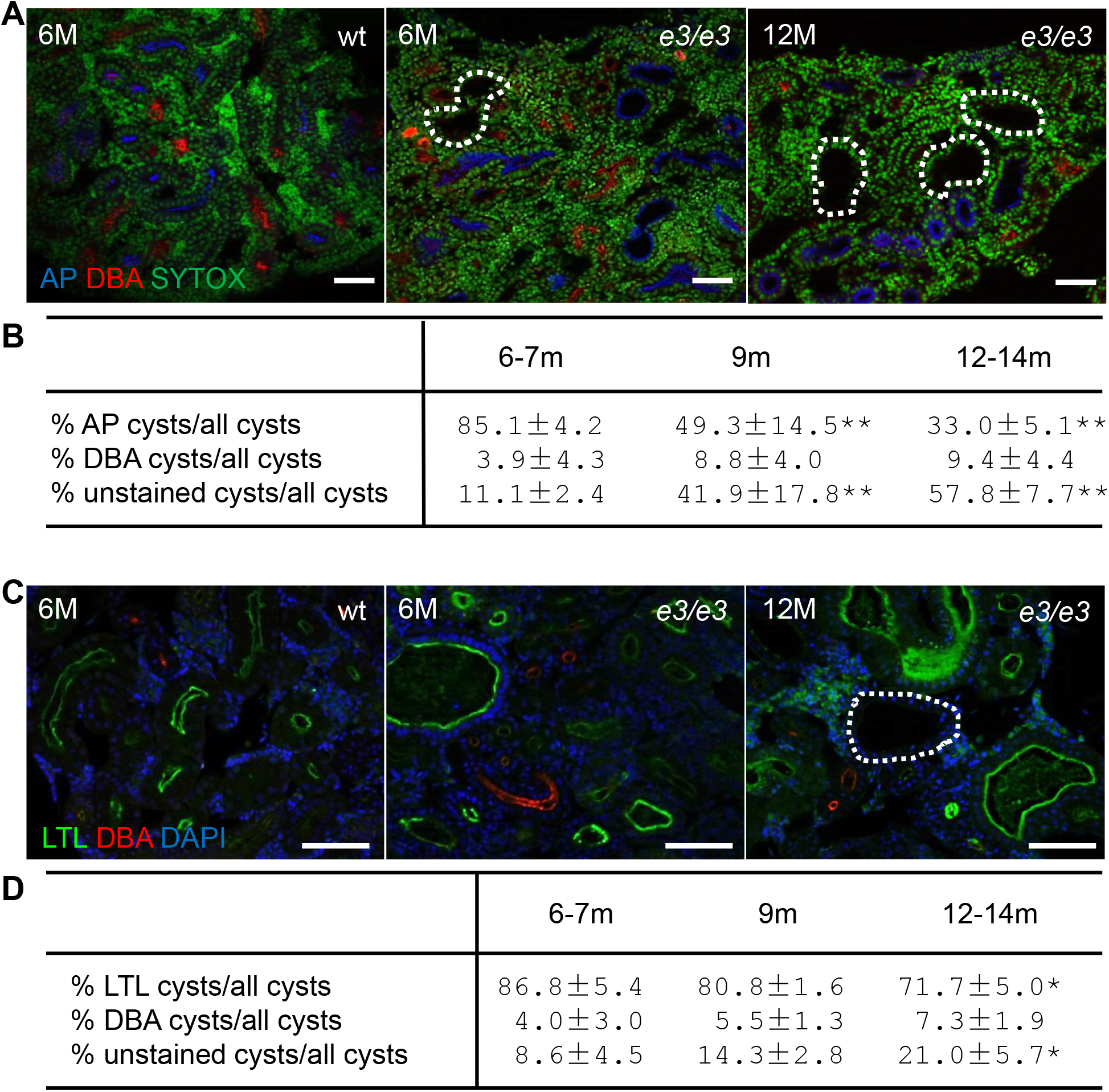
Cystic tubules in *tmem67*^*e3/e3*^ fish are mainly of PT origin at an early stage and CD/dedifferentiated tubule origin later. (**A,B**) Dynamic changes in PT cysts. The tubular origins of cysts were revealed by costaining of frozen sections with AP (blue; labels PTs), DBA (red; labels DTs), and SYTOX (green; labels nuclei) (A). The percentage of segment-specific cysts/total cysts was quantified (B). (**C,D**) Dynamic changes in other cysts. The identities of cystic tubules were indicated by costaining of paraffin sections with LTL (green; labels PTs and major CDs), DBA (red), and DAPI (blue; labels nuclei) (C). The percentage of segment-specific cysts/total cysts was quantified (D). The dashed circle indicates an unstained tubule (A,C). Four to six male fish per genotype at each of the indicated ages were used for renal cyst quantification (B,D). The data are presented as the mean ± s.d. from three independent experiments. *: *P*<0.05; **: *P*<0.01. Scale bar: 20 μm.

One of the characteristics of mammalian PKD is increased proliferation of cyst-lining epithelial cells.^49, 50^ Consistently, we detected more PCNA-positive cells in the *tmem67*^*e3/e3*^ mutants than in those of wild-type fish (Figure 5A,B). Hyperproliferation was observed after cyst formation and was correlated with disease severity (Figures 5B and 7B).

**Figure 5.**
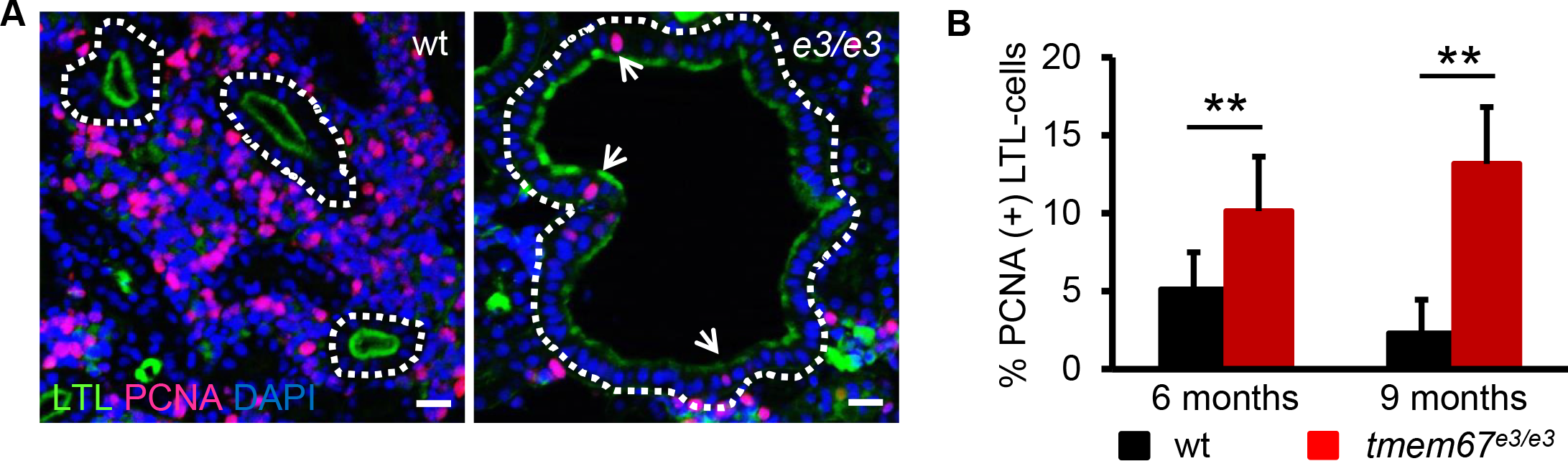
*tmem67*^*e3/e3*^ fish exhibit increased proliferation of cyst-lining epithelial cells. (**A**) Kidney epithelial cell proliferation was assessed by PCNA incorporation. Paraffin sections of 9-month-old fish kidneys were immunostained using a PCNA (red) antibody, and then costained with LTL (green) and DAPI (blue). (**B**) The percentage of PCNA(+) LTL-labeled cells (arrow in A) were quantified in 6- and 9-month-old kidneys. Four to eight male fish per group per time point, 3 sections per kidney, and approximately 200 LTL-labeled cells per section were analyzed. The data are presented as the mean ± s.d. **: *P*<0.01. Scale bar: 20 μm.

### The mesonephric kidneys of *tmem67*^*e3/e3*^ fish have shorter and fewer single cilia but more MCCs than wild-type fish

Because adult zebrafish kidney cilia have not been characterized, we conducted EM analysis. In agreement with the light microscopic observations,^27, 29^ we found that some tubules were lined with thick brush borders, which are indicative of PT segments (Figure 6A,A’), while others had scanty microvilli and could not be clearly distinguished as DTs or CDs (Figure 6B,B’). Cilia bundles were observed in PTs, as noted in other teleost fish, but also in DTs/CDs (Figure 6A,A’,B,B’,G).^51^ Approximately 20-30 cilia measuring 100-150 μm in length were typically clustered together with the distinctive “9+2” motile cilia structure (Figure 6C,G). Single cilia were scarcely detected by SEM (Figure 6H), probably because longitudinal sections were rarely obtained from the thin kidney tissues. In the *tmem67*^*e3/e3*^ kidneys, the epithelial cells in dilated PTs often had loose brush borders (Figure 6D,D’). Clusters of 100-150-μm-long cilia also resided in the PTs and DTs/CDs and retained the 9+2 organization (Figure 6D-F,I). Notably, the mutants appeared to have shorter single cilia than the wild-type fish, although only a few were identified (Figure 6J), and contained more cilia bundles in the PT segment (Figure 6K).

**Figure 6.**
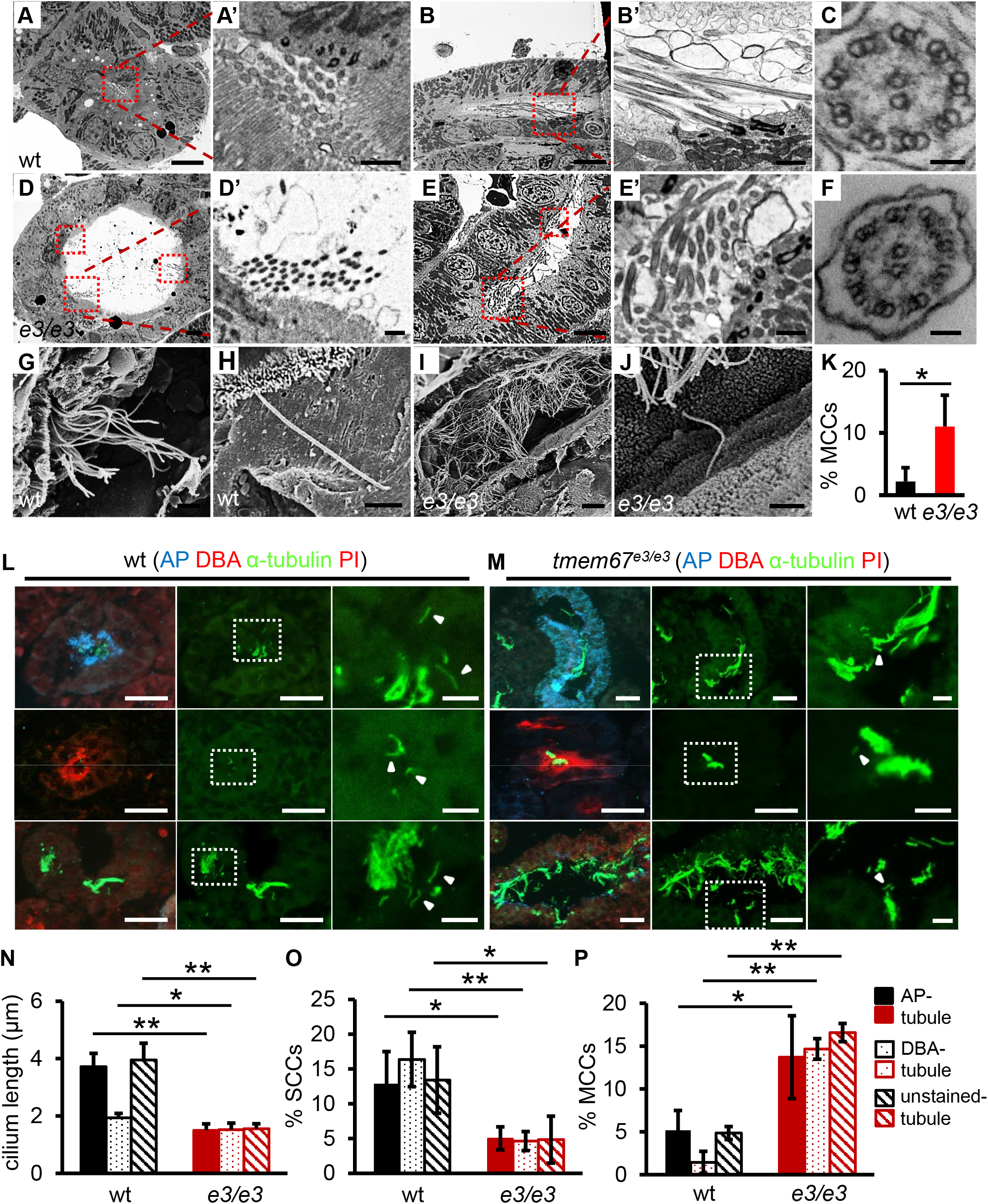
Adult *tmem67*^*e3/e3*^ kidneys have more MCCs and shorter/fewer single cilia than wild-type kidneys. (**A-C**) TEM analysis of cilia in the mesonephric kidneys of wild-type fish. Multicilia bundles (dashed squares) were observed in the PTs (with brush borders) (A) and DTs/CDs (without brush borders) (B), and enlarged (A’,B’). These cilia had typical 9+2 motile cilia structures (C). (**D-F**) TEM analysis of cilia in the mesonephric kidneys of *tmem67*^*e3/e3*^ fish. Multicilia bundles were also observed in the PTs (D,D’) and DTs/CDs (E,E’) and had 9+2 structures (F). (**G-J**) Multicilia bundles (G,I) and single cilia (H,J) in the PT segments of wild-type fish (G,H) and *tmem67*^*e3/e3*^ mutants (I,J) were revealed by SEM analysis. (**K**) The percentage of MCCs in the PT segment was quantified via TEM analysis. Kidneys were collected from 9-month-old male fish, 6 wild-type siblings and 3 mutants, and ~200 PT cells per kidney were scored. (**L,M**) Cilia were examined by α-acetylated tubulin (green) antibody staining of frozen sections. The sections were colabeled with AP (blue), DBA (red), and trace amounts of Propidium Iodide (PI) (red; labels nuclei). Shown from left to right are merged images, cilia-only images (green channel), and enlargement of the boxed area. The arrowhead indicates a single cilium. (**N-P**) The single-cilium lengths in AP-stained, DBA-stained, and unstained tubules were measured (N), the percentages of single-ciliated cells (SCCs) were quantified (O), and the percentages of MCCs were determined (P). Four male fish per genotype were examined at 9 months. A total of 300-800 cells in each segment were counted. The data are presented as the mean ± s.d. *: *P*<0.05; **: *P*<0.01. Scale bars: 5 μm (A,B,D,E,L,M), 1 μm (A’,B’,D’,E’), 50 nm (C,F), 20 μm (G,H,J), and 80 μm (I).

In parallel, we examined single cilia by immunostaining. We noted that single cilia were shorter in DTs than in PTs and AP/DBA(−) tubules, and the latter two had comparable cilium lengths (Figure 6L,N). In *tmem67*^*e3/e3*^ fish, the single cilia were stunted, and the numbers of SCCs were reduced in all segments assessed (Figure 6M-O). Consistent with EM analysis, cilia bundles, as specified by strong α-tubulin staining, were present in all tubular segments and were significantly increased in number in the mutants (Figure 6L,M,P). Together, our data reveal the occurrence of consistent ciliary defects in *tmem67*^*e3/e3*^ embryos and adult fish.

### Ciliary defects are more prominent than cyst development in the mesonephric kidneys of adult *tmem67*^*e3/e3*^ fish

To assess whether pronephric tubule dilation affects mesonephric cyst formation, we raised +EC embryos separately from −EC embryos. The cystic index in +EC fish progressively increased from 4 to 12 months; however, surprisingly, it was lower at 4 months in +EC fish than in −EC fish (Figure 7A). Consistently, epithelial cell hyperproliferation was detected only in −EC kidneys (Figure 7B). Thus, we conclude that cyst development during embryogenesis does not predispose *tmem67* fish to mesonephric cyst formation.

**Figure 7.**
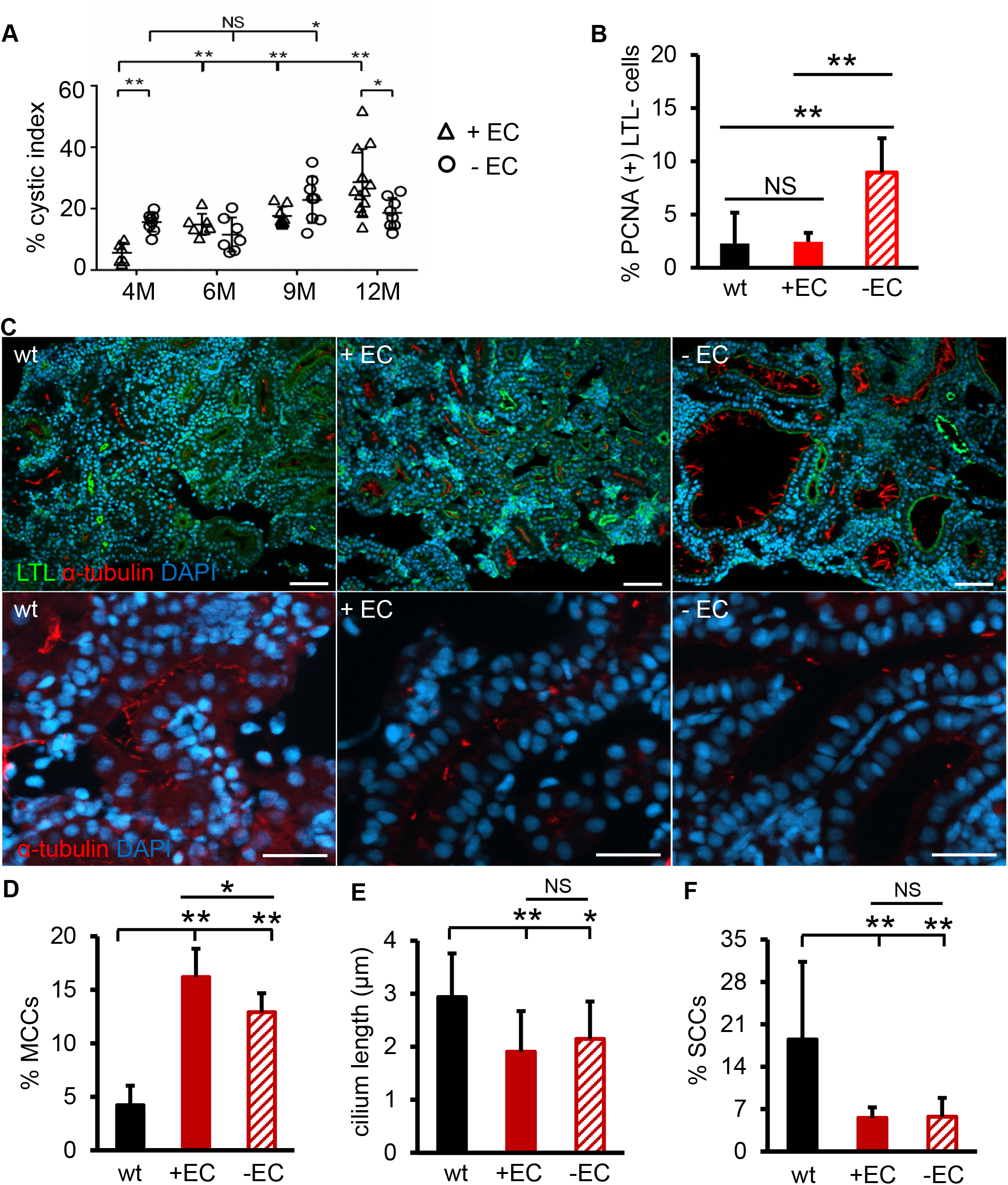
Renal ciliary abnormality precedes mesonephric cyst formation. (**A**) The presence of pronephric cysts did not predispose fish to early mesonephric cystogenesis. Adult *tmem67*^*e3/e3*^ fish that exhibited pronephric cysts (+EC) and those that did not develop pronephric cysts (−EC) were subjected to renal cyst analysis at the indicated ages. The cystic indices quantified from 6-12 male fish per group per time point are shown. (**B**) Cell proliferation in the +EC and −EC fish at 4 months, as indicated by PCNA immunostaining. (**C**) Renal cilia in the +EC and −EC fish at 4 months. Immunostaining of paraffin sections was carried out using a α-acetylated tubulin (red) antibody, and the sections were costained with LTL (green) and DAPI (blue). The upper panels mainly show cilia bundles, and the lower panels were selected to specifically show single cilia. To clearly present single cilia, LTL labeling was removed from the lower panels. (**D-F**) Quantification of the percentages of MCCs (D), single-cilium lengths (E), and percentages of SCCs (F) in the LTL tubules of +EC and −EC male fish at 4 months. Four fish per group and ~250 LTL cells per kidney were examined. The data are presented as the mean ± s.d. *: *P*<0.05; **: *P*<0.01; NS: not statistically significant (*P*>0.05). Scale bar: 20 μm.

Prompted by the significant difference in cyst formation between +EC and −EC fish at 4 months of age, we compared their renal cilia. We found more cilia bundles in the LTL tubules in +EC animals than in −EC animals, although the numbers of MCCs were also substantially increased in the −EC group (Figure 7C,D). On the other hand, the single-cilium length and number were similarly reduced in +EC and −EC fish (Figure 7C,E,F). Taken together, our data suggest that the loss of single cilia is a primary defect of cystogenesis, while MCC expansion might be a later compensatory event.

### mTOR inhibition ameliorates cyst formation, increased proliferation of cyst-lining cells, and ciliary abnormality in *tmem67*^*e3/e3*^ fish

Because the mTOR pathway is often activated and mTOR inhibition is therapeutic in PKD models,^52–54^ we examined mTOR activity in *tmem67*^*e3/e3*^ fish. Indeed, the S6 ribosomal protein was hyperphosphorylated in both +EC embryos and adult mutants (Figure 8A, Supplemental Figure 8A). We also found mTOR activation in *bpck* mice, suggesting a conserved mechanism (Supplemental Figure 9).

**Figure 8.**
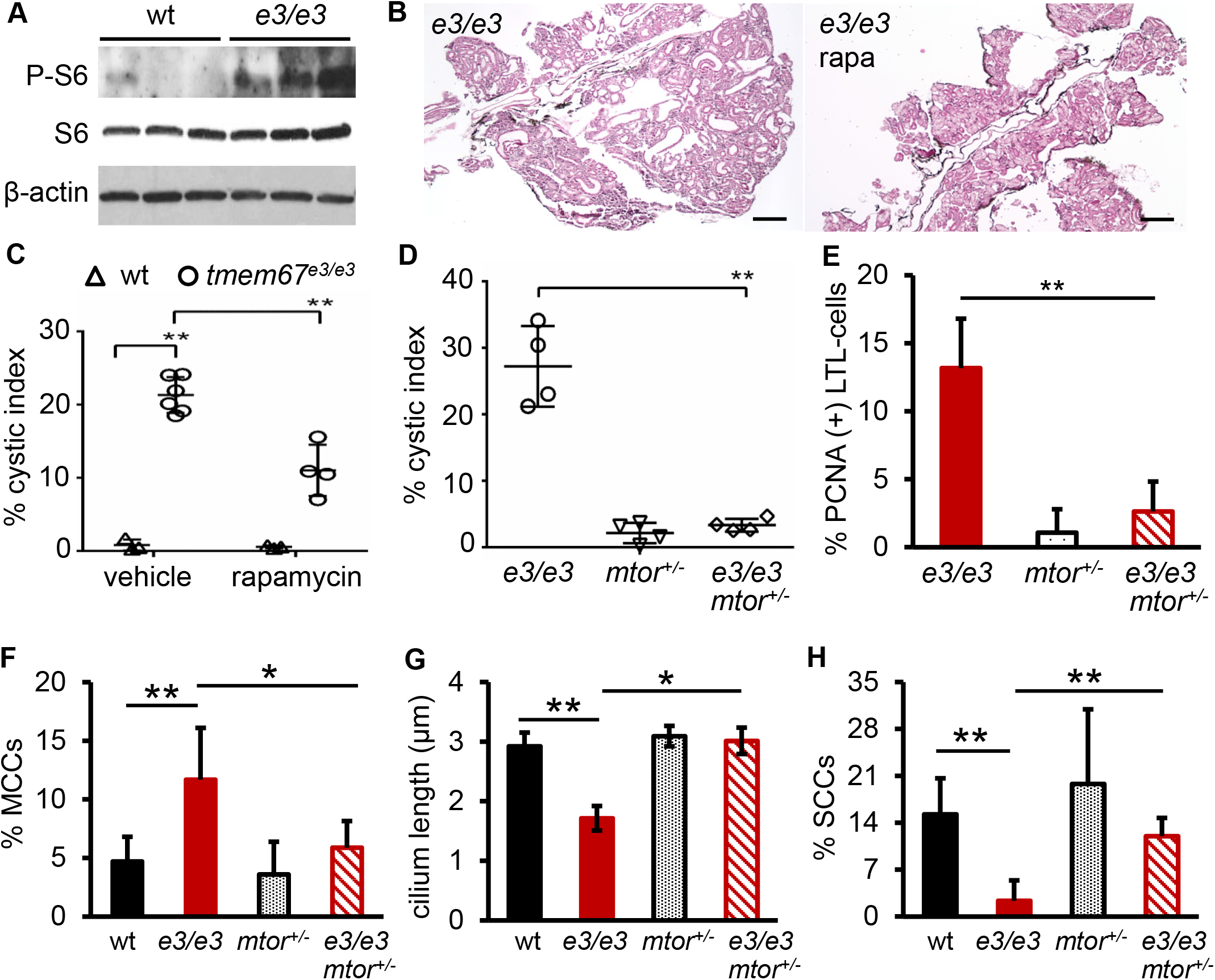
mTOR haploinsufficiency reverses cystogenesis, overproliferation, and ciliary abnormality in *tmem67*^*e3/e3*^ fish. (**A**) Phospho-S6 protein levels were enhanced in the kidneys of 9-month-old *tmem67*^*e3/e3*^ fish. Representative images of three independent experiments are shown. (**B,C**) Rapamycin treatment suppressed cystogenesis in *tmem67*^*e3/e3*^ fish. Rapamycin or vehicle (DMSO) was given to 7-month-old fish by daily gavage for 1 month. Then, renal cyst formation was examined by HE staining of paraffin sections. Representative HE images (B) and the cystic indices (C) are shown. Four to six male fish per group were analyzed. (**D**) *tmem67*^*e3/e3*^;*mtor*^+/−^ double mutants exhibited fewer renal cysts than *tmem67*^*e3/e3*^ single mutants. (**E**) *tmem67*^*e3/e3*^;*mtor*^+/−^ double mutants had fewer PCNA(+) renal epithelial cells than *tmem67*^*e3/e3*^ single mutants. (**F-H**) Ciliary phenotypes in *tmem67*^*e3/e3*^ mutants were rescued in *tmem67*^*e3/e3*^;*mtor*^+/−^ double mutants, as indicated by thepercentages of MCCs (F), single-cilium lengths (G), and percentages of SCCs (H). Four male fish per genotype at 9 months were examined, and all analyses were performed on LTL tubules (D-H). The data are presented as the mean ± s.d. *: *P*<0.05; **: *P*<0.01. Scale bar: 100 μm.

To test whether mTOR inhibition exerts beneficial effects, we treated zebrafish with rapamycin. We found that rapamycin alleviated pronephric cysts in +EC embryos and diminished mesonephric cysts in the adult mutants, without causing apparent defects on wild-type fish (Figure 8B,C, Supplemental Figure 8B). Moreover, genetic inhibition of mTOR signaling with a hypomorphic *mtor*^+/−^ mutant also significantly inhibited cyst growth in the adult fish (Figure 8D).^31^ At the cellular level, we noticed that hyperproliferation of renal epithelial cells was reversed, the numbers of MCCs were normalized, and the single-cilium lengths and percentages of SCCs were restored in *tmem67*^*e3/e3*^;*mtor*^+/−^ double mutants (Figure 8E-H). However, given that short-term rapamycin treatment was unable to alleviate ciliary defects in the embryos (Supplemental Figure 8C,D), the therapeutic benefit of sustained mTOR inhibition likely does not occur via direct regulation of ciliogenesis.

## Discussion

### The adult zebrafish *tmem67* model exhibits key features of mammalian PKD

In this study, we have demonstrated, for the first time, that renal cystogenesis and cilium synthesis can be studied in adult zebrafish, thus establishing a new vertebrate model of PKD that can be used for therapeutic development. Conserved pathogenic processes were noted in adult *tmem67*^*e3/e3*^ fish mesonephric kidneys, including progressive cystogenesis, an age-dependent switch in tubular cyst origin, and increased cell proliferation. Unlike zebrafish embryonic models, the adult model provides a platform for the quantification of cyst size and number. Notably, by following fish with or without pronephric cysts to adulthood, we found that embryonic cysts do not predispose but rather delay mesonephric cystogenesis in young adults. While we cannot rule out the possibility of distinct mechanisms underlying pronephric and mesonephric cyst formation, our observations underscore the additional research opportunities offered by our adult fish model, which can be used to study longitudinal progresses consisting of both compensatory and decompensatory processes.

Compared to MKS3 patients and related rodent models, the adult zebrafish *tmem67* mutants exhibited milder cystic phenotypes. This difference is unlikely an issue related to freshwater species per se, since naturally occurring *pc/glis3* mutants of medaka, another freshwater teleost, develop massive fluid-filled cysts within 6 months.^51, 55^ We hypothesize that this difference can be explained by at least two mechanisms. First, zebrafish have a strong regenerative capacity, not only repairing nephrons (as mammals do) but also forming new ones *de novo*.^56–60^ Because impairments in renal tubule repair after injury have been suggested to trigger cystogenesis,^61, 62^ an elevated regeneration potential would certainly help to slow disease progression. Second, TALEN/CRISPR-mediated mutagenesis has recently been shown to sometimes induce genetic compensation by family members of the targeted gene or other genes in the same biological pathway.^46, 47^ Indeed, transcriptional adaptation of other MKS genes was noted in the *tmem67* kidneys. Thus, identification of critical regeneration/compensation genes or pathways in this fish model might suggest new therapeutic interventions for MKS3 treatment in mammals.

### Role of ciliary abnormality in the zebrafish *tmem67* models

In contrast to the partial penetrance of pronephric cysts, all mutant embryos uniformly exhibited defective single cilia in the distal pronephros, even before cyst formation. The loss of single cilia phenotype also extended to other renal tubular segments of adult fish, including +EC fish at the precyst stage of 4 months. The fact that a lack of cilia preceded cyst development strongly suggests that abnormal ciliogenesis is a primary defect in the *tmem67* model.

On the other hand, multicilia clusters seemed unaffected before 4 dpf. This observation could explain the absence of pronephric cysts in most *tmem67* embryos, because fluid can be propelled out of the pronephros by beating multicilia.^23, 25^ The +EC population likely has additional as-yet-unidentified defects in fluid output. In adult zebrafish kidneys, MCCs were present in all tubular segments, suggesting their roles in normal renal physiology, which are different from those in mammals. MCCs are rarely detected in adult mammals; however, they have been observed in human fetuses, patients with hypercalcemia or nephrotic syndrome, and rodent models of ARPKD.^11, 14, 63–68^ With regard to TMEM67 in particular, more MCCs have been noted in human MKS3 fetuses than in normal fetuses;^11^ in addition, multiple centrosomes and more than one cilium have been detected in *wpk* rat and *Tmem67*-deficient cells *in vitro*, though further investigation is required to determine whether these cells are MCC-like.^11, 14^ Nonetheless, the reappearance of MCCs in diseased kidneys indicates their importance and has been postulated as an adaptive event to relieve local fluid accumulation.^69^ Our data indicating that +EC fish have more MCCs from 4 dpf to 4 months and fewer renal cysts at 4 months than −EC fish provide the first experimental evidence to support this hypothesis.

The zebrafish *tmem67* model recapitulates tissue-specific single cilium loss and MCC induction but not the ciliary elongation phenotype present in some mammalian models.^9, 11, 13, 15^ Cilium length elongation has also been proposed as an adaptive response.^70, 71^ Whether such an adaptive event predominantly presents as cilium lengthening in mammals and MCC expansion in zebrafish due to species-specific expression of ciliogenic factors and/or different susceptibility to cell fate transition warrants further study.

### The adult zebrafish MKS3 model facilitates the identification of candidate therapeutic strategies such as mTOR inhibition

Consistent with findings in PKD patients and various animal models, we noted hyperactivation of the mTOR pathway and beneficial effects of mTOR inhibition in both *tmem67* embryos and adult fish. Because mTOR is also activated in *bpck* mice, mTOR inhibition is anticipated to also be beneficial in rodent MKS3 models. Our results thus support the hypothesis that abnormal mTOR signaling is a common pathological event for PKDs of different etiologies and that mTOR inhibition is a broadly applicable therapeutic strategy.

mTOR signaling has been shown to regulate cilium biogenesis, but the conclusions of the related studies have not always been consistent, and the relationship between mTOR and cilia during cystogenesis remains elusive.^72–78^ We found that long-term mTOR inhibition normalized ciliary abnormalities in adult fish, while short-term inhibition in embryos did not, suggesting an indirect role of mTOR in ciliogenesis. Therefore, we favor the hypothesis that sustained mTOR inhibition exerts therapeutic benefits via a cilia-independent mechanism. Future studies using the adult zebrafish model are anticipated to uncover this mechanism, which could potentially be shared among PKDs of different etiologies, thus facilitating our understanding of the interplay among mTOR signaling, ciliogenesis, and cystogenesis.

## Author Contributions

X.L. and X.X. designed the study; P.Z., Q.Q. and X.L. carried out experiments; P.Z. and X.L. analyzed the data and made the figures. X.L., X.X. and P.C.H. drafted and revised the paper; all authors approved the final version of the manuscript.

## Acknowledgements and Financial Disclosures

We thank Bingquan Huang and Scott I. Gamb of the Mayo Clinic Electron Microscopy Core Facility for expert TEM and SEM assistance, Amanda C. Leightner for kindly providing us with mouse kidney samples. This work was supported by the Mayo Translational Polycystic Kidney Disease Center (MTPC) Pilot and Feasibility grant (NIDDK DK90728) (to X.L.) and the Mayo Foundation for Medical Education and Research (to X.X.).

Disclosures: None

## Table of Contents for the Supplemental material

Supplemental Figure 1. *tmem67* is expressed in multiple tissues of the adult zebrafish

Supplemental Figure 2. Small percentage of *tmem67*^*e3/e3*^ embryos develops hydrocephalous

Supplemental Figure 3. *tmem67*^*e3/e3*^ mutants do not show exon skipping events

Supplemental Figure 4. Ciliary and MCC defects in *tmem67*^*e3/e3*^ embryos at 4 dpf

Supplemental Figure 5. Cilium lengths are not significantly altered in the Kupffer’s vesicle and neural tubes

Supplemental Figure 6. Anterior migration of the pronephric epithelial cells and convolution of the proximal tubules are defective in *tmem67*^*e3/e3*^ embryos

Supplemental Figure 7. The expression of other MKS genes in *tmem67*^*e3/e3*^ kidneys

Supplemental Figure 8. The effect of rapamycin on pronephric cyst formation and cilium biosynthesis in *tmem67*^*e3/e3*^ embryos

Supplemental Figure 9. mTOR is activated in *bpck* mice

