## Supplemental Materials for "*mtor* Haploinsufficiency Ameliorates Renal Cyst Formation in Adult Zebrafish *tmem67* Mutants"

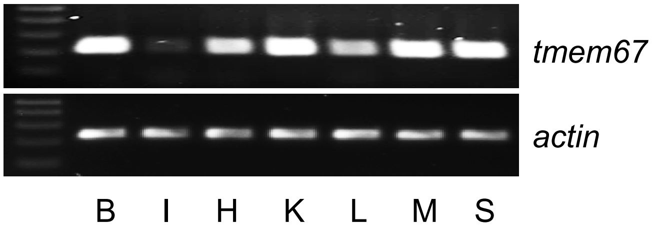


**Supplemental Figure 1.** ***tmem67* is expressed in multiple tissues of the adult zebrafish**
*tmem67* transcripts were detected in the brain (B), heart (H), kidney (K), liver (L), muscle (M), spleen (S), and weakly in the intestine (I) via RT-PCR.

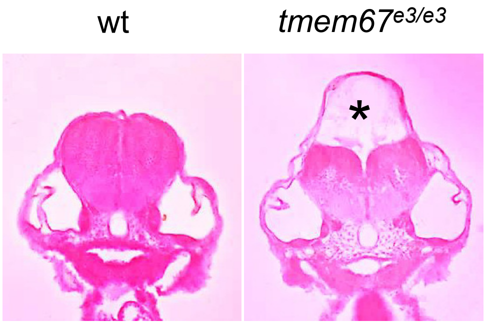


**Supplemental Figure 2. Small percentage of *tmem67*$^{e3/e3}$ embryos develops hydrocephalous**
Embryos at 3 dpf were assessed for the presence of hydrocephalous by HE staining. Three out of 85 *tmem67*$^{e3/e3}$ fish showed hydrocephalous (asterisks) while 0 out of 79 wild-type embryos did.

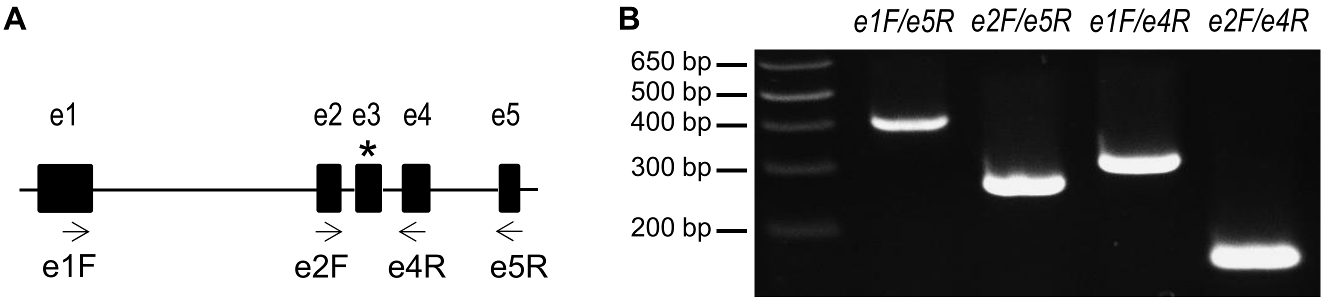


**Supplemental Figure 3. *tmem67*$^{e3/e3}$ mutants do not show exon skipping events**
(**A**) Schematic diagram showing exons surrounding the targeted exon3 of *tmem67* gene and primer binding sites. (**B**) RT-PCR amplification of cDNAs from the *tmem67*$^{e3/e3}$ embryos. Without alternative splicing events, PCR products are predicted to be 422 bp using e1F/e5R, 276 bp using e2F/e5R, 316 bp using e1F/e4R, and 170 bp using e2F/e4R, respectively. *tmem67-e1F*: TTCTCCATATCATTTCGACAGC; *tmem67-e2F*: GGTCAGTAATGGGGTGTCTATC; *tmem67-e4R*: AGGGATTGCCATTCCCATC; *tmem67-e5R*: GAGTTAATGAATGAGTCCCCAC.

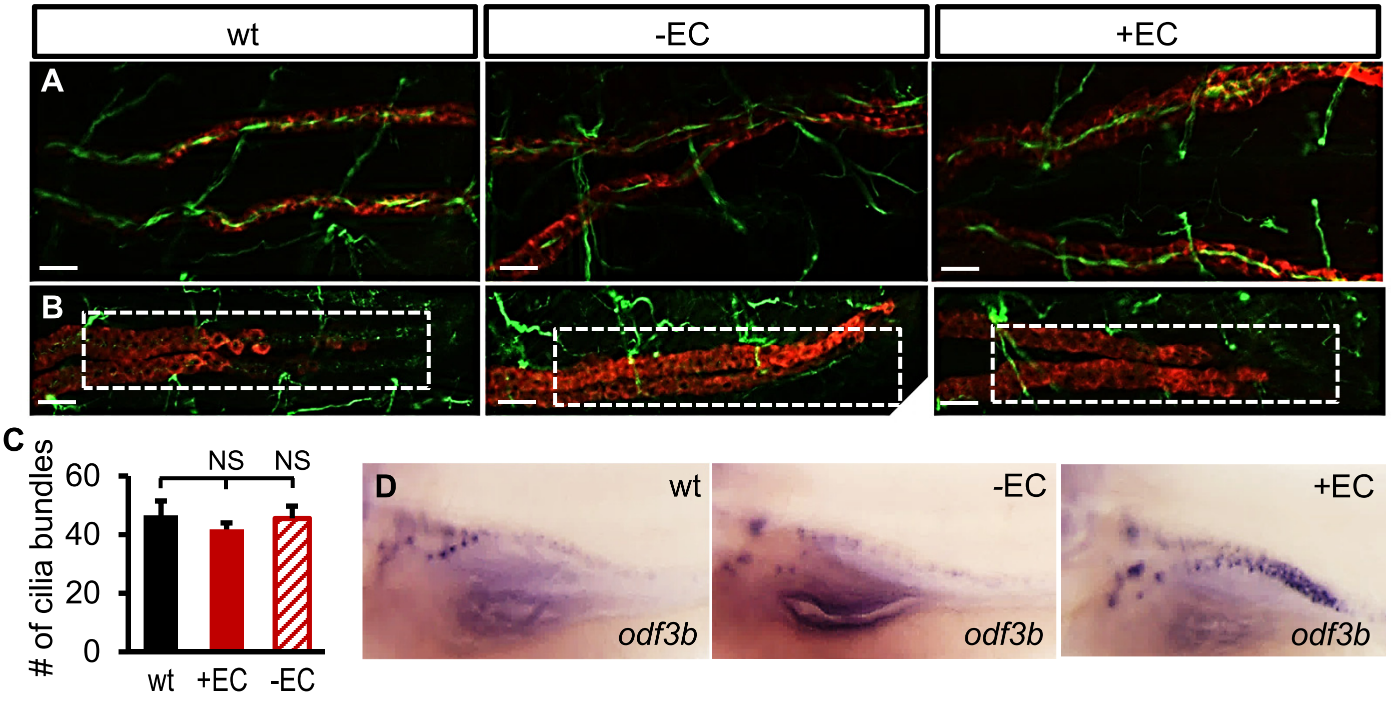


**Supplemental Figure 4. Ciliary and MCC defects in *tmem67*$^{e3/e3}$ embryos at 4 dpf**
(**A,B**) Multicilia bundles (A) and distal single cilia (B) of the pronephros were shown by co-immunostaining using antibodies against a-acetylated tubulin (green) and $Na^+/K^+$ ATPase (α6F, red). **(C)** Quantification of multicilia bundles. (**D**) MCCs were indicated by *in situ* hybridization using riboprobe against *odf3b*. A total of 10-12 embryos per group per experiment were examined. Scale bar: 10 µm.

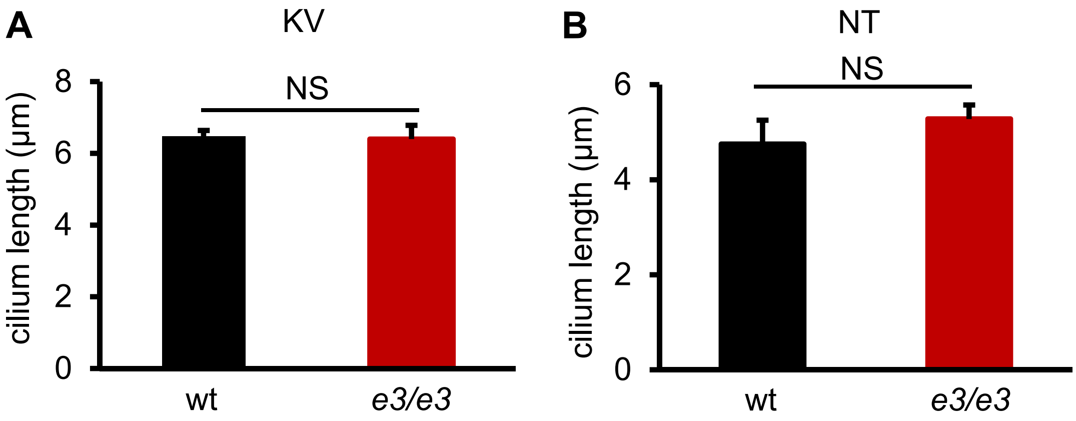


**Supplemental Figure 5. Cilium lengths are not significantly altered in the Kupffer's vesicle and neural tubes**

Embryos were collected at 10 somites (A) and 26 hpf (B) for cilia length analysis in the Kupffer's vesicle (KV, A) and neural tubes (NT, B), respectively. Eight to 12 embryos per group were analyzed. NS: not statistically significant.

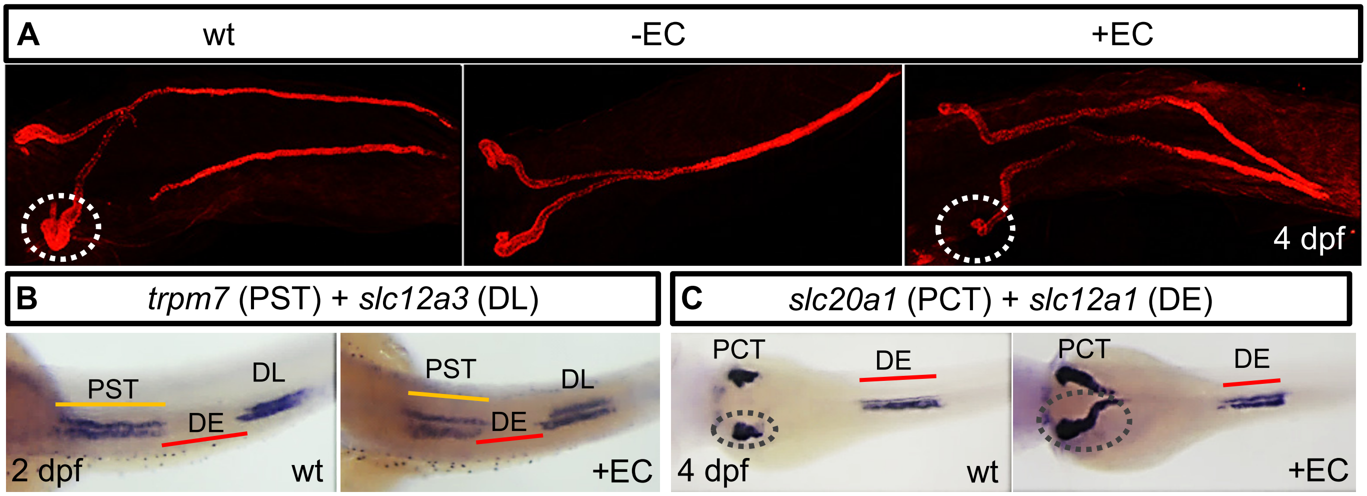


**Supplemental Figure 6. Anterior migration of the pronephric epithelial cells and convolution of the proximal tubules are defective in *tmem67*$^{e3/e3}$ embryos**

(**A**) Lack of convolution of the proximal tubule (dashed circles) was particularly seen in the +EC embryos (4 out of 4). Shown are whole-mount immunostaining using a6F antibody. (**B,C**) +EC embryos had shorter PST and DE segments at 2 dpf (B, 7 out of 9) and less convoluted PCT fragments (C, dashed circles, 11 out of 12). Shown are *in situ* hybridization using riboprobes against *slc20a1a, trpm7*, *slc12a1*, and *slc12a3*, which are markers for PCT, PST, DE, and DL segments, respectively.

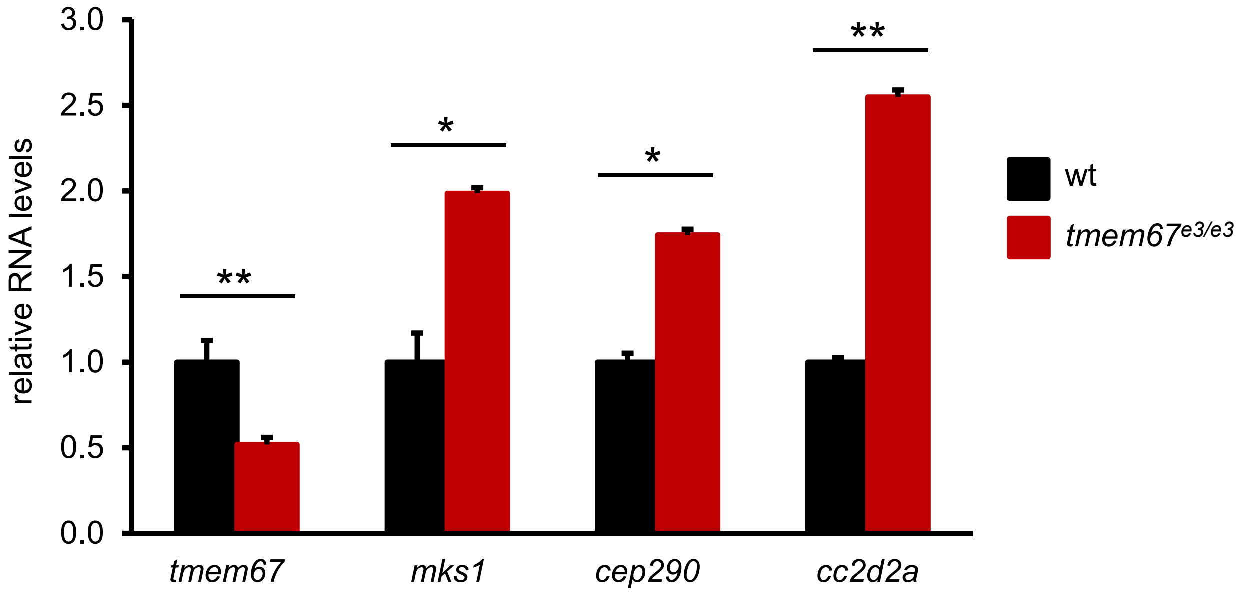


**Supplemental Figure 7. The expression of other MKS genes in *tmem67*$^{e3/e3}$ kidneys**
Kidneys from *tmem67*$^{e3/e3}$ fish and wild type siblings were collected at 9 months, and subjected to RNA extraction, cDNA synthesis and Q-PCR analysis of MKS genes, including *tmem67* (*MKS3*), *mks1*, *cep290* (*MKS4*), and *cc2d2a* (*MKS6*). Primers used for Q-PCR are as follows: *tmem67-F*: AGAATCTGTGTGGCCAAAGG, *tmem67-R*: CAATGACAGAGCCCAGGAAT; *mks1-F*: TGAGGGCTATGGCTATTTGG, *mks1-R*: CAGTGGTCTGAGTGCGAAAG; *cep290-F*: ACCACTGACGAAGGTCTTGC, *cep290-R*: CAGCTCTTGGGTCAGTTTCC; *cc2d2a-F*: AACCTCTGGAAGCCGTTTTT, *cc2d2a-R*: GGCAGAAACTGGCGTAATGT. Three mutants and 3 wild type fish were used, and gene expression was normalized by *gapdh*. *: $P$<0.05. **: $P$<0.01.

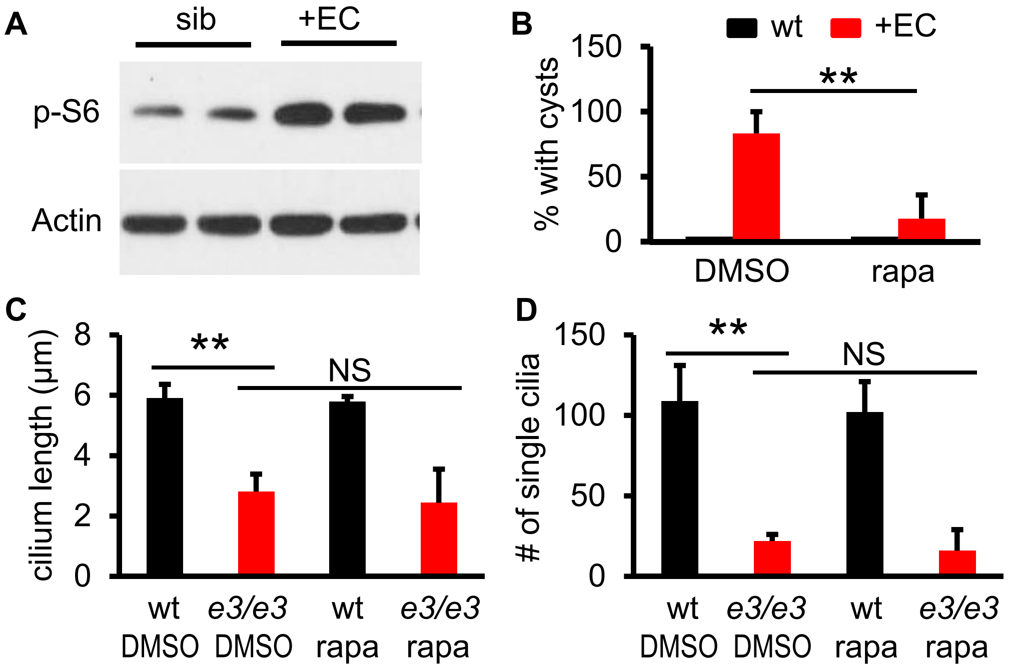


**Supplemental Figure 8. The effect of rapamycin on pronephric cyst formation and cilium biosynthesis in *tmem67*$^{e3/e3}$ embryos**

(**A**) Phosphorylation of S6 protein was increased in +EC embryos at 5 dpf. Shown are representative images of three independent experiments. (**B**) Rapamycin treatment reversed pronephric cyst formation. +EC or wild-type embryos were incubated with rapamycin (400 nM) or DMSO at 56 hpf for 16 hours. The percentages of embryos with kidney cysts were analyzed by HE staining of JB-4 sections. (**C, D**) Rapamycin did not restore distal single cilium length and number. *e3/e3* and wild-type siblings were incubated with rapamycin at 56 hpf for 16 hours, and distal single cilia length and number were quantified following co-immunostaining using antibodies against a-acetylated tubulin and $Na^+/K^+$ ATPase (α6F). Twelve to 16 embryos per group were analyzed (B-D). **: $P<0.01$. NS: not statistically significant.

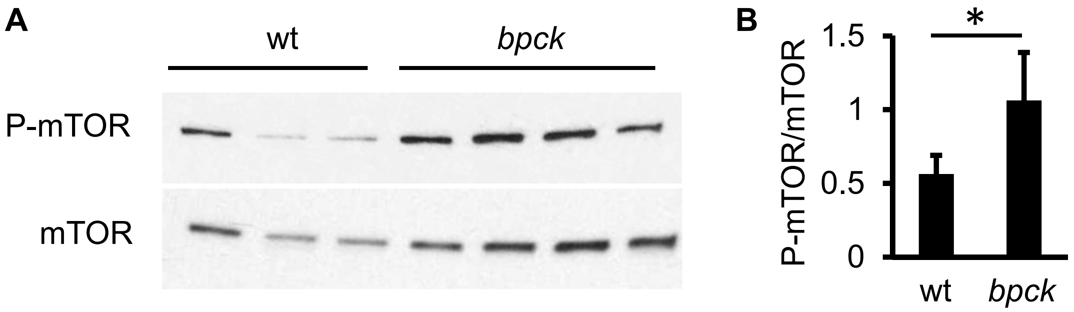


**Supplemental Figure 9. mTOR is activated in *bpck* mice**
Kidneys from 3-week-old wild-type or *bpck* mice were collected. Phosphorylation pf mTOR was analyzed by western blotting (A) and quantified (B). *: $P<0.05$.
